# Lexical Access in Comprehension vs. Production: Spatiotemporal Localization of Semantic Facilitation and Interference

**DOI:** 10.1101/449157

**Authors:** Julien Dirani, Liina Pylkkänen

**Author notes:** **Corresponding author**: Julien Dirani, NYU Abu Dhabi, Saadiyat Island, Abu Dhabi, 129188.

## Abstract

Humans understand words faster when they are preceded by semantically related words. This facilitation is thought to result from spreading activation between words with similar meanings. Interestingly, in language production, semantic relatedness often has the opposite effect: in object naming for example, a related prior word delays the naming time of the current object. This could be due to competition during conceptual search or later interference at the motor preparation stage. However, no study has systematically compared the facilitory and inhibitory effects and thus their neurobiological relationship is unknown. We contrasted maximally parallel production and comprehension tasks during magnetoencephalography and found that in comprehension (specifically word reading), semantic relatedness modulated activity in the left middle STG at 180-335ms, consistent with prior findings on the spatiotemporal localization of lexical access. In contrast, a semantic interference pattern for the production task (object naming) occurred in a post-lexical time-window at 395-485ms in left posterior insular cortex, consistent with post-lexical motor preparation. Thus, our data show that semantic priming during comprehension and interference during production are not two sides of the same coin but rather they clearly dissociate in space and time, consistent with a lexical account for comprehension and a post-lexical one for production.

**Significance statement:** The processing of semantically related words has been a central tool for understanding the organization of the mental lexicon. One striking observation is that semantic relatedness tends to be facilitory in comprehension but inhibitory in language production, perhaps because only production involves a conceptual search through semantically related candidates. The neurobiology of this contrast is not understood. Our magnetoencephalography results demonstrate that the facilitory pattern is first observed in classic left temporal lexical access regions at ~200ms, whereas the inhibitory pattern occurs later and in the insular cortex. These findings show that the two effects do not co-localize in space or time and suggest that the inhibitory effects in production stem from a late motor preparation stage.

## INTRODUCTION

Human language is both an action and a perception system: in language production, we choose a meaning and transform it to generate a motor output, while in comprehension, we perceive linguistic input and map it onto semantic representations. In principle, it is possible to conceive of production and comprehension as mostly using the same processing route in opposite directions, but the extent to which this is actually true is far from understood.

At the lexical level, it is well established that word comprehension is faster when the word is preceded by another semantically related word, or prime. This effect is explained as automatic activation of the prime that spreads to the representation of the second word (Neely, 2012). Interestingly, this effect is reversed in a production task like object-naming, where semantically related primes slow down naming time, an effect known as semantic interference (Lupker, 1979). Both the semantic facilitation and the interference literatures are considerably developed, yet they remain separate. Here, we used the spatio-temporal resolution of magnetoencephalography (MEG) to systematically compare the effects of semantic relatedness in language comprehension and production.

The locus of semantic interference in production is highly debated, with some hypotheses placing it at the word retrieval (i.e. lexical) level and others at later, post-lexical stages. On the lexical account, the effect is caused by competition for selection between the activated representations of the prime and target, and consequently, lexical selection is achieved via competition (Bloem & La Heij, 2003; Levelt, Roelofs, & Meyer, 1999). However, more recent studies have proposed that the effect is post-lexical, at the level of articulatory programs. The Response Exclusion Hypothesis (Janssen, Schirm, Mahon, & Caramazza, 2008; Mahon, Costa, Peterson, Vargas, & Caramazza, 2007) assumes that language production involves a single-channel output buffer to which visually presented words have privileged access over names of images (Mahon et al., 2007). Before the name of the image can be produced, this buffer would have to be cleared of the representation of the prime; a process regulated by semantic information (Glaser & Glaser, 1989; La Heij, 1988; Lupker, 1979), giving rise to the interference effect.

fMRI studies have shown that both semantic interference in production and facilitation in comprehension affect the STG, the MTG, and the ACC. Other frontal regions are also involved, with the IFG showing semantic priming in comprehension and the OFC interference in production (de Zubicaray, Wilson, McMahon, & Muthiah, 2001; Kotz, Cappa, von Cramon, & Friederici, 2002; Maess, Friederici, Damian, Meyer, & Levelt, 2002; Matsumoto, Iidaka, Haneda, Okada, & Sadato, 2005; Rissman, Eliassen, & Blumstein, 2003; Rossell, Price, & Nobre, 2003). Regarding timing, facilitation in comprehension has been observed at 200-500ms (Holcomb & Anderson, 1993; Kiefer & Spitzer, 2000) and interference in production at 200-300ms after stimulus onset using EEG (Aristei, Melinger, & Abdel Rahman, 2011), and slightly earlier in MEG, at 150-225ms (Maess et al., 2002). It seems, then, that priming effects in comprehension and production may share some brain correlates, but it is not clear exactly where and when they overlap.

To clearly delineate the spatiotemporal relationship between semantic facilitation in comprehension and inhibition in production, we measured MEG during highly parallel comprehension and production tasks. If priming effects in both tasks localize in middle temporal cortex around 300ms or earlier, this would strongly conform to a shared lexical-level origin. In contrast, if the interference effect in production is manifested later than 400ms after picture onset, a motor-preparation account of this effect is more likely. Crucially, we included identical primes to discriminate between facilitory and inhibitory neural patterns. Since identical prime-target pairs (table-table) are maximally related and expected to elicit robust facilitory repetition priming, we considered a pattern facilitory if the semantically related condition patterned between the unrelated and identity conditions. In contrast, in an inhibitory pattern, the semantically related condition should diverge from both the identity and unrelated conditions, neither of which should show similarity-based interference.

**Figure 1.**
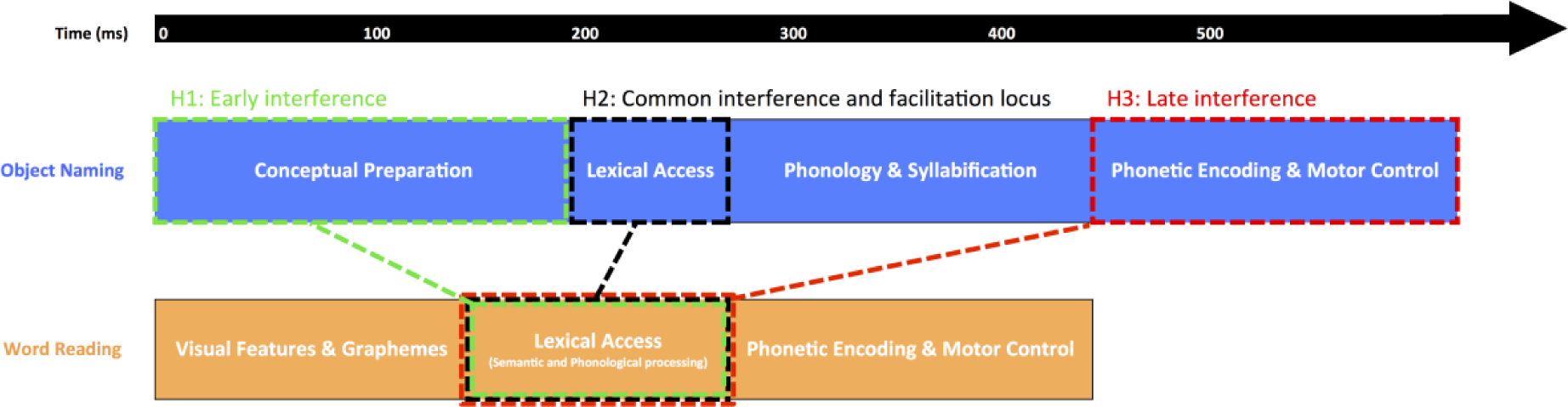
Three contrasting hypotheses (H1, H2, H3) regarding the localization of semantic interference in Object Naming. Semantic and phonological processing is depicted as occurring in parallel to lexical access in Word Reading.

## METHODS

### Participants

Thirty right-handed native English speakers were recruited to take part in the study. Two participants were excluded due to excessive artifacts that resulted in >25% of rejected trials, and 3 participants were rejected due to equipment failure, leaving 25 good participants (14 Female, M=22.67, SD=5.55). All participants had normal or corrected-to-normal vision and reported no history of neurological or language disorders. The study received ethical approval from the Institutional Review Board at New York University Abu Dhabi.

### Experimental design

The experiment consisted of a comprehension and a production task. The challenge was designing maximally parallel tasks to allow adequate comparison across them. Specifically, production includes a motor component which is absent in comprehension. We resolved this by using overt Word Reading for the comprehension task and Object Naming for production. Thus, we were able to manipulate conceptual search, which is the key difference between production and comprehension, while maximizing the similarity between tasks, since reading a word does not require a conceptual preparation (Indefrey & Levelt, 2004). This is in agreement with computational models of reading such as the Dual Route Cascading model which describe reading and reading aloud equivalently (Coltheart, Rastle, Perry, Langdon, & Ziegler, 2001). Further, brain activity associated with reading aloud and comprehension tasks (such as lexical decision) have been linked to shared word recognition and lexical retrieval areas (Carreiras, Mechelli, Estévez, & Price, 2007). There is therefore sufficient support for the assumption that here, overt Word Reading does indeed operationalize a comprehension task.

The targets that had to be named consisted of line drawings for Object Naming, and lowercase words for Word Reading. In both tasks, three levels of primes were manipulated: First, identical primes (Ident), that were the repetition of the target for words, and the name of the object for images. We included this condition to provide us with a clear facilitation effect for both tasks, which would allow us to interpret the remaining effects in comparison. This is crucial for the brain data, where an increase or decrease in activation is not always straightforward to interpret. We also used semantically related primes (Semrel), which were words that belonged to the same semantic category of the targets. Categorically related primes have been shown to reliably induce interference effects in Object Naming. We therefore only used this type of semantic relation in order to guarantee to observe the interference priming effect that we aim to compare to facilitation in the comprehension task. Finally, unrelated primes (Unrel) were words that differed from the target in all aspects (visual, phonology, and semantics). All primes were in capital letters while all target words were in lowercase to control for purely visual priming. We also manipulated the stimulus onset asynchrony (SOA) at 4 different intervals: 150, 200, 250, and 300 milliseconds (ms). These SOAs were in the range that showed reliable interference effects in Object Naming (Heij, Dirkx, & Kramer, 1990; Sailor, Brooks, Bruening, Seiger-Gardner, & Guterman, 2009), but it was not clear how priming in Word Reading would be affected. The four chosen SOAs allowed us to explore whether priming in Word Reading could at any point turn into an interference effect. Importantly, the potential interaction of SOA with Prime Type and Task could provide us with additional insight as to the timing of priming effects.

Stimuli were presented using Psychopy 1.84.2 (Peirce, 2007) on a screen positioned above the participants’ heads while they laid back on a bed in the magnetically shielded room of the MEG. Each trial started with the presentation of a fixation cross that appeared for 300ms, followed by a blank screen for 300ms. Next, the prime appeared for 100ms, followed by a blank screen. The duration of the prime was held constant, but the blank screen following it varied to create an SOA of 150, 200, 250, or 300ms depending on the condition being presented. Finally, the target stayed on screen until the participants named it (Figure 2). Responses were recorded with a microphone positioned near the participant’s mouth.

**Figure 2.**
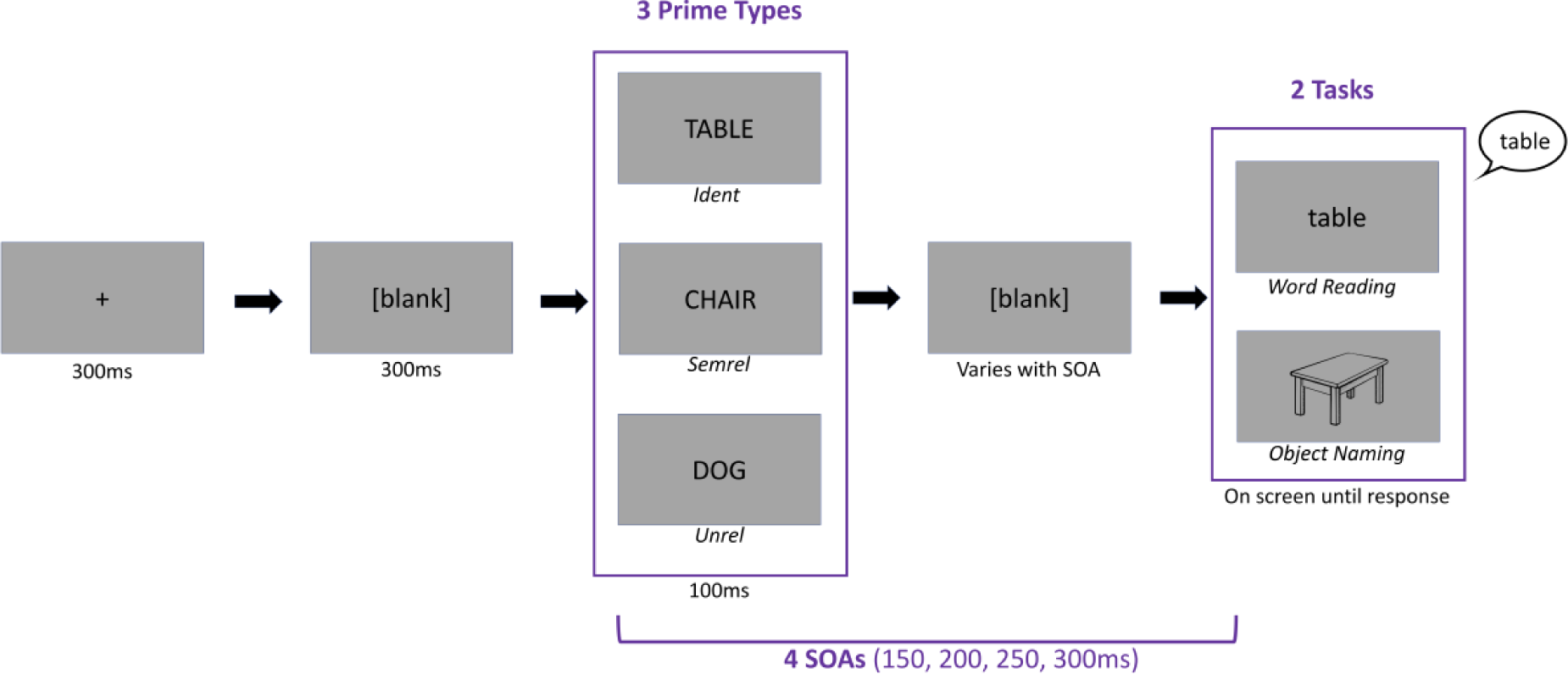
Trial structure and experimental design.

### Stimuli

The lists of all primes and the list of targets were English nouns in their root form, balanced for length and frequency across all lists. After the stimuli generation was done, 50 participants rated the semantic relation between the unrelated and semantically related primes and the targets via the Amazon Mechanical Turk platform (www.mturk.com). They were instructed to rate how much they thought the words belonged to the same category using a Likert scale ranging from 1 to 5. Any prime-target pairs that had an average rating between 2.5 and 3.5 were excluded from the stimuli, creating 2 distinct groupings of semantically related and unrelated prime-targets (Semrel pairs: M=4.31, SD=0.32; Unrel pairs: M=1.25, SD=0.19).

The stimuli consisted of 82 Sets. Within each Set, there was one common target that was repeated 6 times: 3 times as an image and 3 times as a word. There were unique Semrel and Unrel primes for each of the target types. The Ident prime was repeated twice, once with each target type. Since we also wanted to manipulate SOA at four intervals (150, 200, 250, 300), we would have ideally proceeded in one of the following two way, but each was problematic: either every Set would be presented with all of the four SOAs, but then the items would repeat excessively (potentially allowing participants to anticipate upcoming targets) or, we would create completely unique Sets of stimuli for each SOA, balanced on relevant characteristics. However, it turned out to be unfeasible to generate that many unique Sets of prime-target pairs, controlled in all the necessary ways. As a compromise, we opted to present each one of our Sets of stimuli (i.e. 6 prime-target pairs, with a shared target), twice. In order to control for anticipation and predictability, we created 2 versions of each prime type while trying to minimize the semantic distance between the 2. In other words, each Set was presented twice, changing the versions of the primes in each repetition. The result was that subjects saw each target 12 times (6 times as an image, and 6 times as a word), and each unique prime twice (once before the image, and once before the word), with the exception of the identity prime that was seen 4 times. In order to avoid confounding any effects of SOAs with effects created by specific items, it was necessary to avoid consistently pairing specific SOAs with specific Sets. That is, we had to counterbalance the pairing of SOAs with Sets across subjects. This was done by first arbitrarily splitting the 82 Sets into two lists of 41 Sets. Each list of Sets (A and B) was then paired with two SOAs, thus presenting each Set twice (as previously stated). This pairing was counterbalanced across every six participants in order to ensure that a specific pairing of SOA to item did not have an undue influence on the group-level results. Further, in order to control for the order of repetitions of targets within subjects, each of the 24 cells of the design was assigned to a block number following a Latin-squared method. Thus, the block number corresponded to the order of trials in the experiment. This was done to ensure that within each subject, the number of times that a given condition (e.g. Unrelated, Object Naming, 150ms SOA) appeared earlier in the experiment than another condition (e.g. Semrel, Word Reading, 200ms SOA) was equal, pairwise across all conditions in the experiment. However, since the total number of cells in the design was 24 (6 prime-target pairs x 4 SOAs), while the total number of items in a was List 42, it was not possible to fully cross all of the conditions with block number (which would require 48 Sets per List, 96 in total). Thus, we distributed each of the conditions across block number in a manner that was as close to uniform as possible. Within the resulting 12 blocks, trial order was randomized, which conserved the Latin-squared order over the whole trials.

### MEG acquisition and processing

Continuous MEG was recorded with a 208-channel axial gradiometer system (Kanazawa Institute of Technology) at a sampling rate of 1000 Hz with an online band-pass filter of 0.1- 200Hz. The raw data was noise-reduced with the continuously adjusted least-squares method (Adachi, Shimogawara, Higuchi, Haruta, & Ochiai, 2001) using the MEG Laboratory software 2.004A (Yokogawa Electric and Eagle Technology Corp., Japan). All the following preprocessing was done using the MNE-Python 0.14 (Gramfort et al., 2014) and Eelbrain 0.25.2 (Brodbeck, 2017) packages. The data was first converted to .fif format. After visual inspection of the data, bad channels were excluded, and the data was low-pass filtered offline at 40Hz. An Independent Component Analysis was then fitted to the data using the “fastica” method, selecting components by 95 cumulative percentage of explained variance. Components related to eye-blinks, heartbeats, saccades, and dead channels where then rejected manually. Epochs from −400 to 700ms from stimulus onset were extracted. Since the SOA manipulation changed the timing of the pre-stimulus baselines relative to the target onset, baseline correction was done differently with each SOA making sure all baselines fell during the 100ms before the onset of the. Epochs exceeding a maximum peak-to-peak threshold of ±2000 femto-tesla were removed automatically, and the remaining epochs were scanned for eye-blink artifacts and were removed accordingly. Finally, wrong responses, responses in which participants stuttered, and responses faster than 300ms and slower than 2000ms were excluded from the analysis. All the remaining good epochs were down-sampled by 5, so that the sampling rate became 200Hz, and were then averaged by condition to form the evoked responses.

Each subject’s head-shape was created using an optical FastSCAN scanner (Polhemus) and was co-registered with the FreeSurfer (http://surfer.nmr.mgh.harvard.edu/) average brain. In order to execute a better co-registration, the average brain was scaled using 3-dimensional axes to match each subject’s head-shape. The source space was defined as a dipole grid on the white matter surface using the topology of a recursively subdivided icosahedron (“ico-4” option). Only sources in the left hemisphere were included, and were defined using the PALS-B12 atlas (Van Essen, 2005). A separate inverse solution was then computed for each subject with the evoked responses, using the forward solution as well as the noise covariance matrix computed from the respective 100ms baselines of each condition. For each subject, the noise covariance matrix was estimated using the best estimator out of the three methods ‘shrunk’, ‘diagonal_fixed’, and ‘empirical’, based on log-likelihood and cross-validation on unseen data (Engemann & Gramfort, 2015). For each source location, minimum norm current estimates were computed using 3 orthogonal dipoles, resulting in a 3D vector. Only the lengths of the vectors were retained, resulting in orientation-free source estimations. The resulting estimates were noise-normalized at each source using an SNR regularization factor of 3. This resulted in noise-normalized statistical parametric maps (SPMs), which were then converted to dynamic maps (dSPMs), and provided information about the statistical reliability of the estimated signal at each source (Dale et al., 2000). Finally, source activity was morphed to the FreeSurfer average source space in order to be comparable across subjects.

### Statistical analysis

The initial statistical analysis was based on a mass univariate analysis with spatiotemporal cluster-based permutation tests (Holmes, Blair, Watson, & Ford, 1996; Maris & Oostenveld, 2007). Average source estimates for each condition and for each subject were used in the analysis. The F value of a 2 × 3 × 4 repeated-measures ANOVA was computed for each source at each time point in the full left hemisphere and limited to the 100-600ms timewindow. This F-map was thresholded at an F-value corresponding to an uncorrected p-value of 0.01. Clusters were formed based on direct adjacency in space and time, with the restrictions that they contain a minimum of 10 sources and last at least 10ms. The sum of all F-values (ΣF) was computed for each resulting cluster. This procedure was then repeated 10,000 times, each time with a random permutation of the data, by shuffling condition labels within subjects. For each permutation, the largest of the ΣF was saved to create a non-parametric permutation distribution. The Monte Carlo p-value was computed for each cluster in the original F-map as the proportion of random permutations in which the observed ΣF was larger than the values from the permutation distribution. We retained clusters whose Monte Carlo p-value was smaller or equal to 0.05.

A secondary analysis was performed in order to unpack the patterns of priming effects within each Task. The same cluster-based permutation test described above was performed, sub-setting the data by Task and thus using a 3×4 repeated-measures ANOVA (3 Prime Types, 4 SOAs). The Monte Carlo p-value threshold was corrected to 0.025 in order to account for the multiple comparisons across the 2 tasks.

Significant clusters were plotted as time-courses as well as bar graphs showing their average dSPM value. Sources included in the cluster are plotted on the Fsaverage brain with the average F-values for the time-window of the significant cluster. In all plots, time 0 represented the onset of the target. Concerning results for the effect of Prime Type, we only reported clusters showing a semantic priming pattern. That is, we only presented clusters where the Semrel and Unrel conditions showed distinct timecourses that separate from each other. The reason is that pure Ident priming effects do not directly address our hypothesis, since the Ident condition was only included as a baseline for interpreting semantic priming effects.

Voice onset reaction times (RTs) were analyzed with a linear mixed-effect model using the LmerTest package (Kuznetsova, Brockhoff, & Christensen, 2017) in R (Team, 2013). As with the MEG data, wrong responses, responses in which participants stuttered, and responses faster than 300ms and slower than 2000ms were excluded from the analysis. The initial model included all main effects of Prime Type, Task, SOA, all 2-way interactions, and the 3-way interaction as fixed effects. Random intercepts were used for subjects and items. In order to test for the significance of the predictors, we performed a sequential decomposition of the contributions of the fixed-effects using the ANOVA function from the LmerTest package, using type III hypothesis test. For each predictor, an F-test and its corresponding *p* value were estimated using Satterthwaite’s method (Giesbrecht & Burns, 1985; Hrong-Tai Fai & Cornelius, 1996). Post-hoc pairwise comparisons of significant effects were done using differences of least square means corrected for multiple comparisons using the Tukey method, with Satterthwaite’s estimation for degrees of freedom. The final model was then retrieved with backwards elimination of non-significant effects.

## RESULTS

### Behavioral data

We found a main effect of Task (F(1, 23382) = 5585.43, *p* < .001), with longer RTs observed for Object Naming (M = 817.63, SD = 211.25) compared to Word Reading (M = 679.04, SD = 156.10). There was also a main effect of Prime Type (F(2, 23378) = 1053.39, *p* < .001) showing that the Ident condition was the fastest (M = 688.25, SD = 172.55), followed by Unrel (M = 773.51, SD = 194.98), and finally Semrel (M = 779.79, SD = 210.62) (all *p* < .001). Further, we found an interaction between Task and Prime Type (F(2,23377) = 326.16, *p* < .001), which showed that, in Object Naming, RTs were shorter for Unrel (M = 855.76, SD = 200.16) compared to Semrel primes (M = 876.48, SD = 219.71), illustrating the predicted semantic interference effect. In contrast, for Word Reading, RTs were shorter for Semrel (M = 690.57, SD = 155.46) compared to Unrel primes (M = 694.91, SD = 153.06), although this effect was not significant (p = 0.21). The Ident priming condition revealed the fastest RTs in both Tasks. The main effect of SOA was a reliable predictor of RTs (F(3,14703) = 56.21, *p* < .001), and also interacted with the effect of Task (F(3, 23378) = 2.8817, *p* < .05). Within each Task, RTs got shorter as SOAs got longer, with the exception of SOAs 250 and 300 in the Object Naming Task where RTs did not significantly differ (*p* = .20)

The final model that was obtained with backwards elimination of non-significant effects is presented below. Items and subjects were included as random factors.

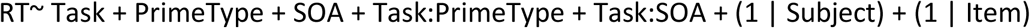

**Figure 3.**
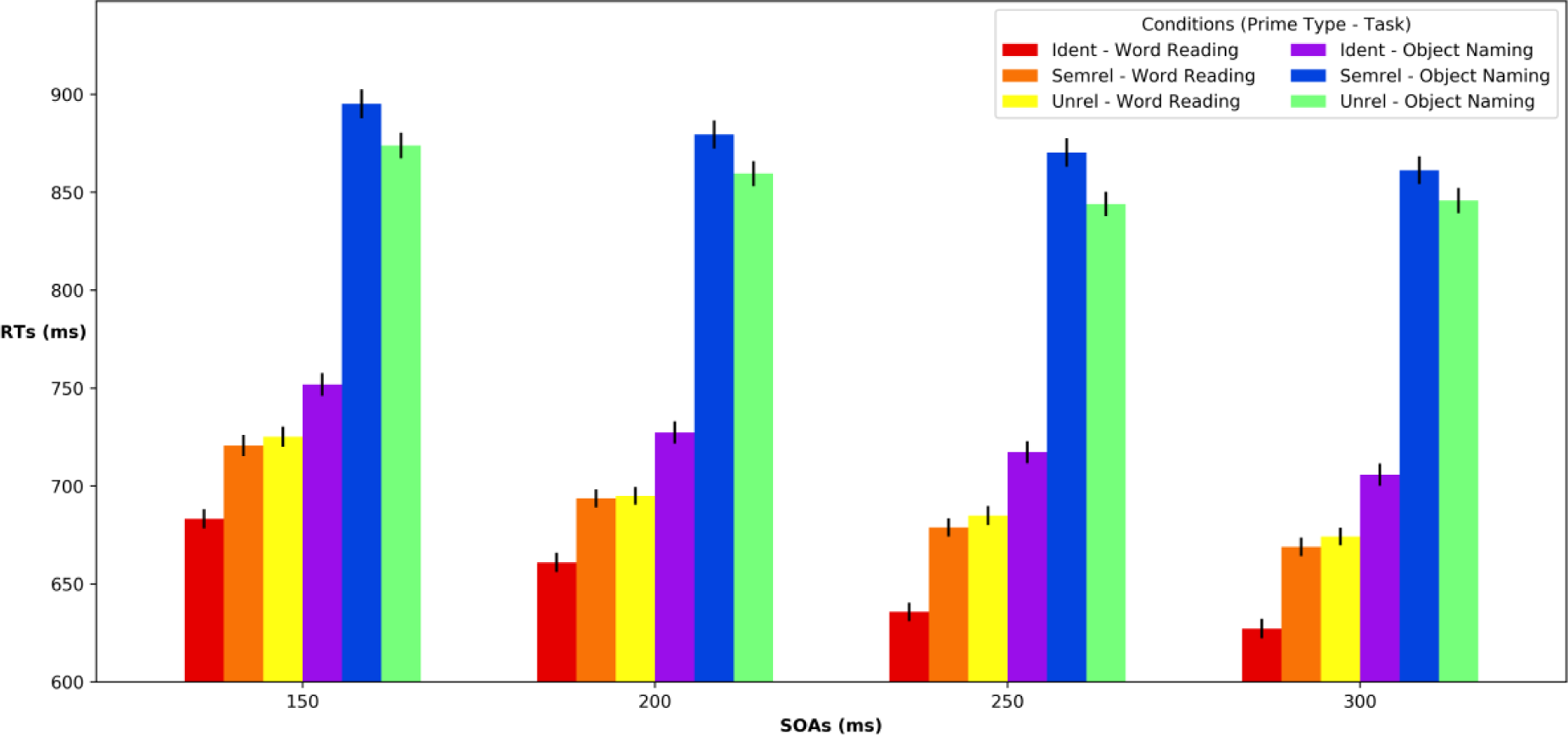
Behavioral reaction times (i.e., utterance onset times) across all conditions. A stable effect of Prime Type was observed in both Word Reading (warm colors) and Object Naming (cool colors) across all SOAs.

### MEG data

#### Omnibus Analysis

The omnibus cluster-based permutation test revealed a robust, widespread main effect of task. The biggest spatial cluster expanded on most of the left hemisphere (93.72%, 2401 sources) and lasted for the full analysis time-window (100-600ms, *p* < .001) (Figure 4). In addition, the timecourse of the two tasks showed two drastically different patterns. This indicates that Word Reading and Object Naming were associated with strikingly different neural signatures across the better part of the left hemisphere.

**Figure 4.**
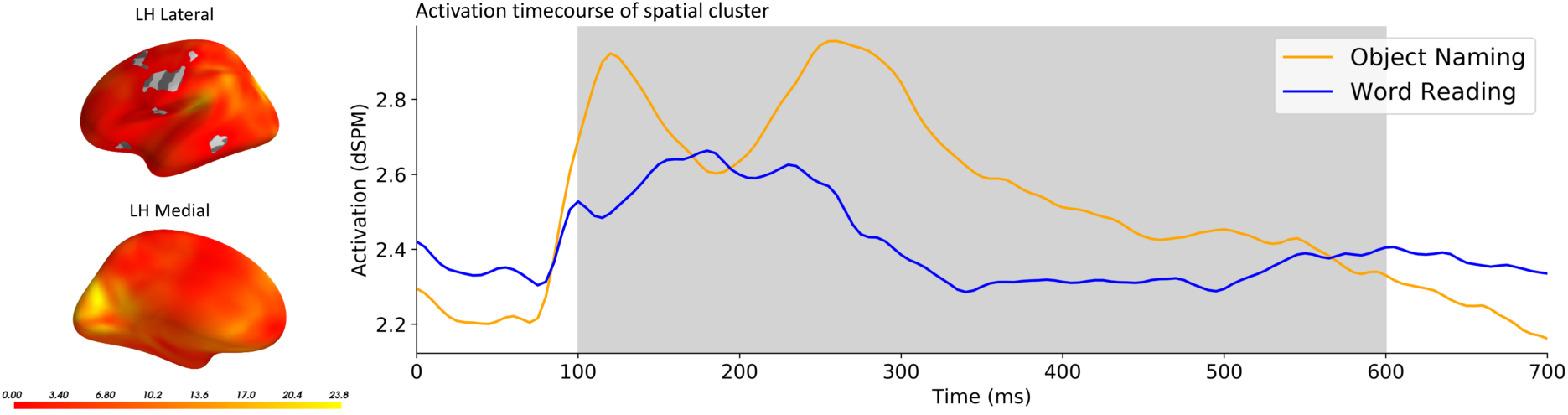
A wide-spread main effect of Task extending through the entire analysis time window and almost the entire left hemisphere (as well as the right hemisphere, as observed in an additional posthoc test; *p* < 0.05, corr.). The large effect of task motivated our within task analyses, to achieve greater sensitivity to observe priming effects.

We also found a spatio-temporal cluster for the main effect of Prime Type (Figure 5) showing a priming effect localized to the middle STG between 175ms and 370ms (p<.001) in which activation increases stepwise as semantic distance increases. This pattern followed that of the behavioral results in which RTs were shortest for Ident primes followed by Semrel primes, and finally Unrel primes.

**Figure 5.**
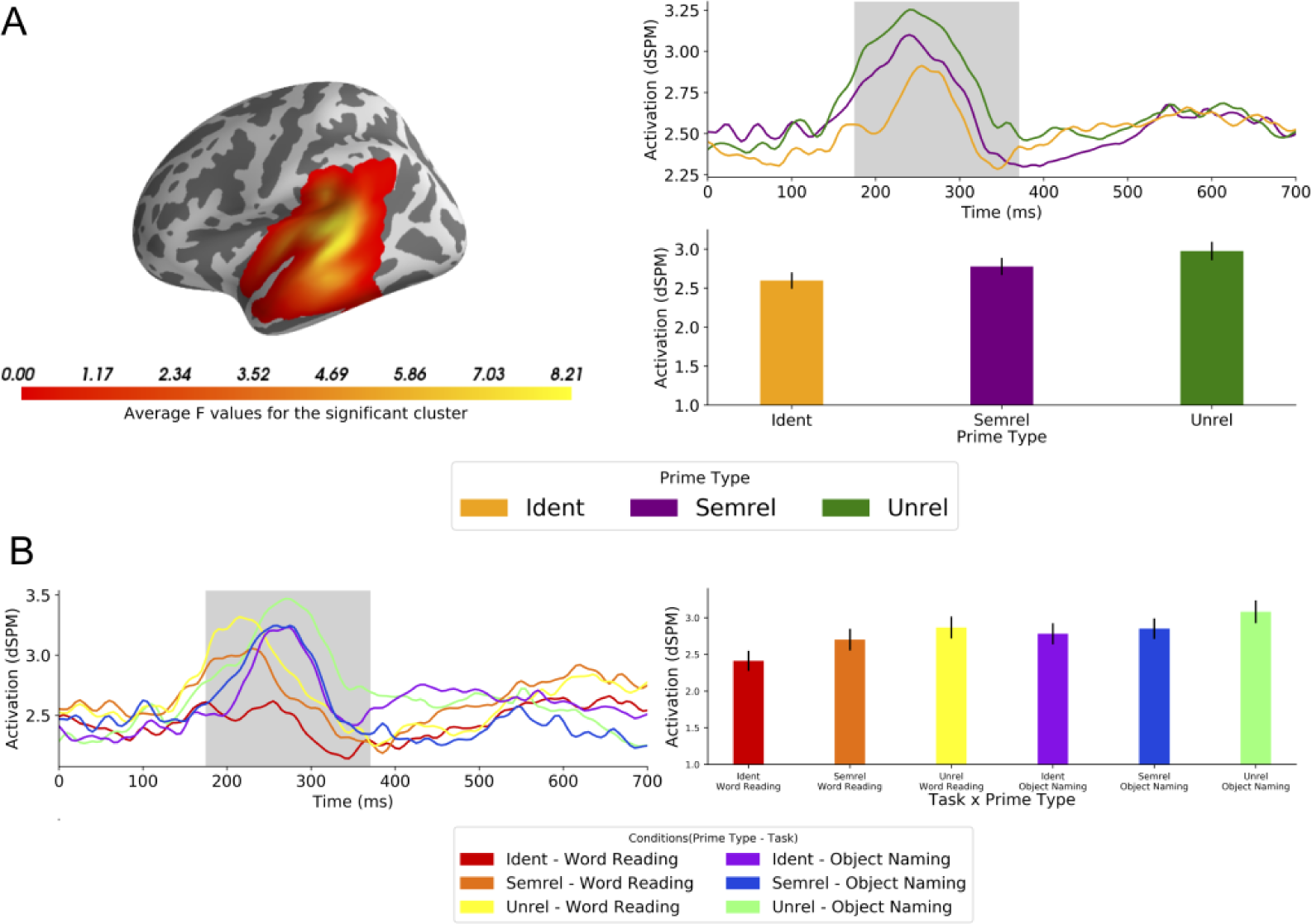
Main effect of Prime Type in the full across tasks analysis. (A) Spatial extent of the significant cluster (right) and its time course averaged across tasks. Bar graph shows mean amplitudes of the significant (shaded) temporal cluster across priming conditions. (B) Same data split by task, revealing a clearer step-wise amplitude reduction for Word Reading than for Object Naming. (Grey shading on time courses indicates *p* < 0.05, corr.).

The effect of Prime Type did, however, interact with the effect of Task (Figure 6), indicating that the priming pattern described above might be a generalization that is not necessarily representative of the priming patterns within each Task. For Object Naming, the priming pattern was in line with the behavioral results, with the highest activation for the Semrel primes, followed by the Unrel primes, and finally the Ident primes. For the Word Reading task, the activation of the Semrel condition appeared higher than that of Unrel and Ident priming conditions, however, the latter 2 had similar activation levels.

**Figure 6.**
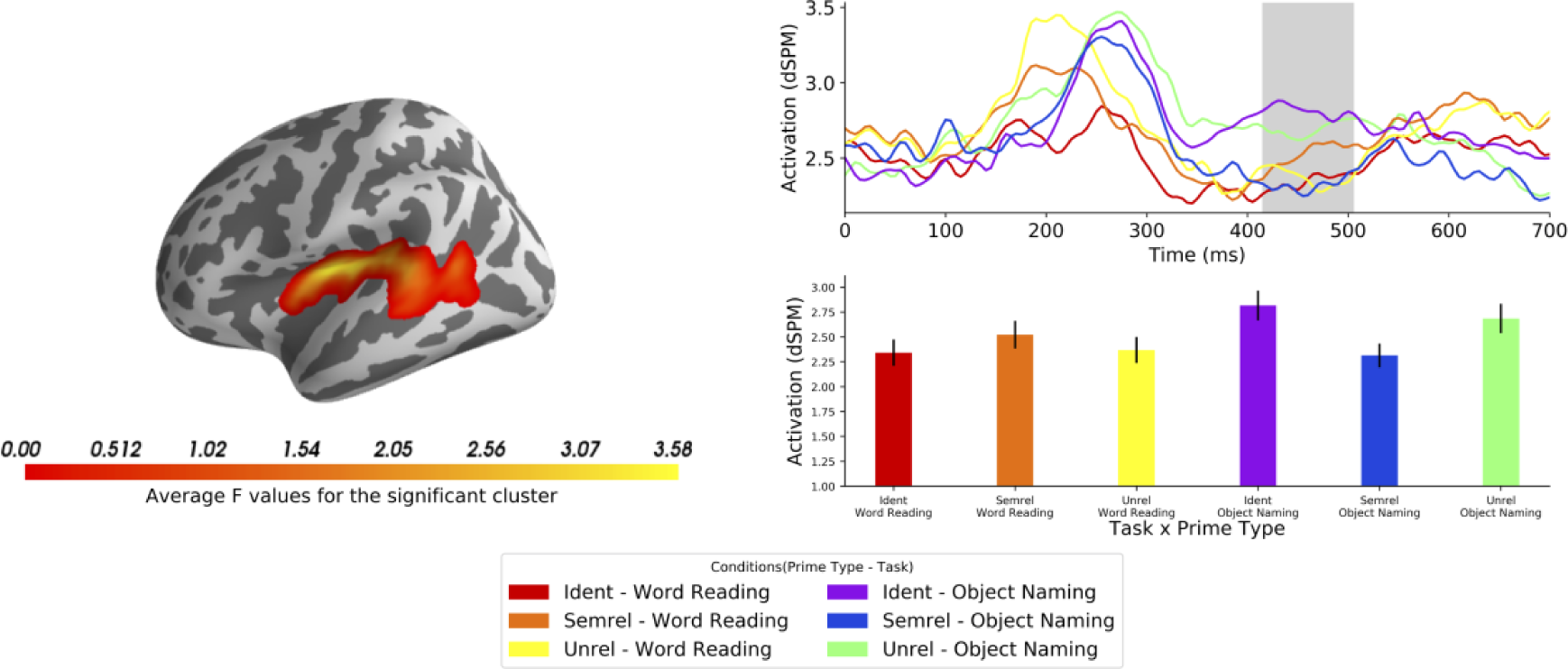
Interaction cluster between Task and Prime Type in the full across tasks analysis, showing an amplitude reduction for the semantically related condition in the Object Naming Task only (*p* Ã 0.05, corr.).

Finally, we found that SOA modulated brain responses at 6 different spatio-temporal clusters. The largest cluster contained 518 sources located in frontal areas as well as anterior medial temporal areas (*p*<.001). This cluster showed a stepwise increase in activation as SOAs got longer, which could be representative of top-down inhibitory processes associated with a suppression of the prime, leading to faster naming times as SOAs got longer. This pattern is compatible with the behavioral data; however, it was not consistent across all of the remaining 5 clusters.

#### Analysis Within Task

Our primary goal was to unpack the neural signatures of semantic priming for Object Naming and Word Reading. Since the main effect of Task was so dramatic, with 93.72% of the left hemisphere modulated by Task for the whole analysis time window, we opted for a second analysis within Task. The motivation was to exclude the large effect of Task in order to have a better understanding of semantic priming within each task. Further, plotting the main effect of prime type within task revealed that the main pattern that was observed (Figure 5.A) does not apply in Object Naming (see Figure 5.B), implying that the effect might be driven by Word Reading alone. The second analysis within task allowed us to directly address this.

With this second analysis, we were able to observe 2 distinct priming patterns, one for each task (Figure 7). For the Object Naming task, we found a late cluster at 395-485ms (*p*<.001), with its center located on the insular cortex with sources reaching the anterior STG. The insular cortex has been associated with motor aspects of speech production, specifically coordination of articulation (Ackermann & Riecker, 2004; Dronkers, 1996; Wise, Greene, Büchel, & Scott, 1999). In addition, the late timing of the effect corresponds with theoretical timings of phonetic encoding and motor preparation for speech (Indefrey, 2011; Indefrey & Levelt, 2004). We observed that the Ident and Semrel conditions separated in opposite directions from the Unrel condition, with higher activation for Ident and lower for Semrel. Facilitation priming in production tasks is expected to induce an increase in activation (Blanco-Elorrieta, Ferreira, Del Prato, & Pylkkänen, 2018), but here we observe a decrease in activation for the Semrel condition compared to the Unrel condition and, importantly, an increase in activation with Ident primes which, overall, represents a semantic interference pattern. This pattern is in line with RT results where Ident primes elicited the fastest responses, followed by Unrel, and finally Semrel primes. Crucially, the priming pattern observed was different in the Word Reading task, where we found an early facilitation priming pattern between 180-335ms (*p*<.001) at the middle STG and expanding to the middle MTG and the posterior part of the insular cortex. We observed a stepwise increase in activation as semantic distance increased, which is typical of facilitatory priming (e.g. Bentin, McCarthy, & Wood, 1985; Holcomb, 1988). This pattern is once again in line with the observed RTs. This cluster appeared to be very similar to the cluster in Figure 5.A, which indicates that the stepwise priming effect found in the first analysis was likely driven by the Word Reading task alone; especially since the second analysis did not find a similar pattern in Object Naming.

Finally, there was a main effect of SOA for both tasks. In Object Naming, we found 4 clusters modulated by SOA, all occurring early, before 200ms. All of the clusters seemed to exhibit the pattern that was observed in the behavioral data, with a stepwise increase in activation, as SOAs got longer. Once again, an increase in activation for Object Naming was associated with shorter naming times. In Word Reading, the posterior part of the insular cortex appeared to be modulated by SOA at 180-210ms. The activations at this cluster did not seem to follow a straightforward pattern. Here, main effects of SOA do not directly address our neuroscience of language questions, but the interaction of SOA with the other main effects of our design could bring insight to the timing of sub-processes of language production and comprehension. For example, if semantic interference effects only occur with short SOAs, we could imply an early locus of the effect. Unexpectedly, in neither of our analyses did the effect of SOA interact with Prime Type. It appeared that the influence of the prime on Word Reading and Object Naming is constant from 150 to 300ms after target onset (since SOA was manipulated for those times). This is supported by the behavioral data that did not show an interaction effect between SOA and Prime Type. We therefore conclude that on average, our priming results generalize across all the SOAs that we tested. Post-hoc visual inspection of the data indeed confirmed that the priming effects found for each task still hold within SOAs. Nevertheless, since there is evidence that shows that SOA manipulation can change the size, and even the direction of priming, specifically in Object Naming (e.g. Heij et al., 1990; Xavier Alario, Segui, & Ferrand, 2000), it is likely that SOAs longer than 400ms might turn out to interact with Prime Type and/or Task. Since the effect of SOA did not modulate that of Prime Type, and the main effect of SOA does not directly address our research questions, we chose not to further discuss its implications. (See supplementary materials for brain clusters associated with the main effects of SOA).

**Figure 7.**
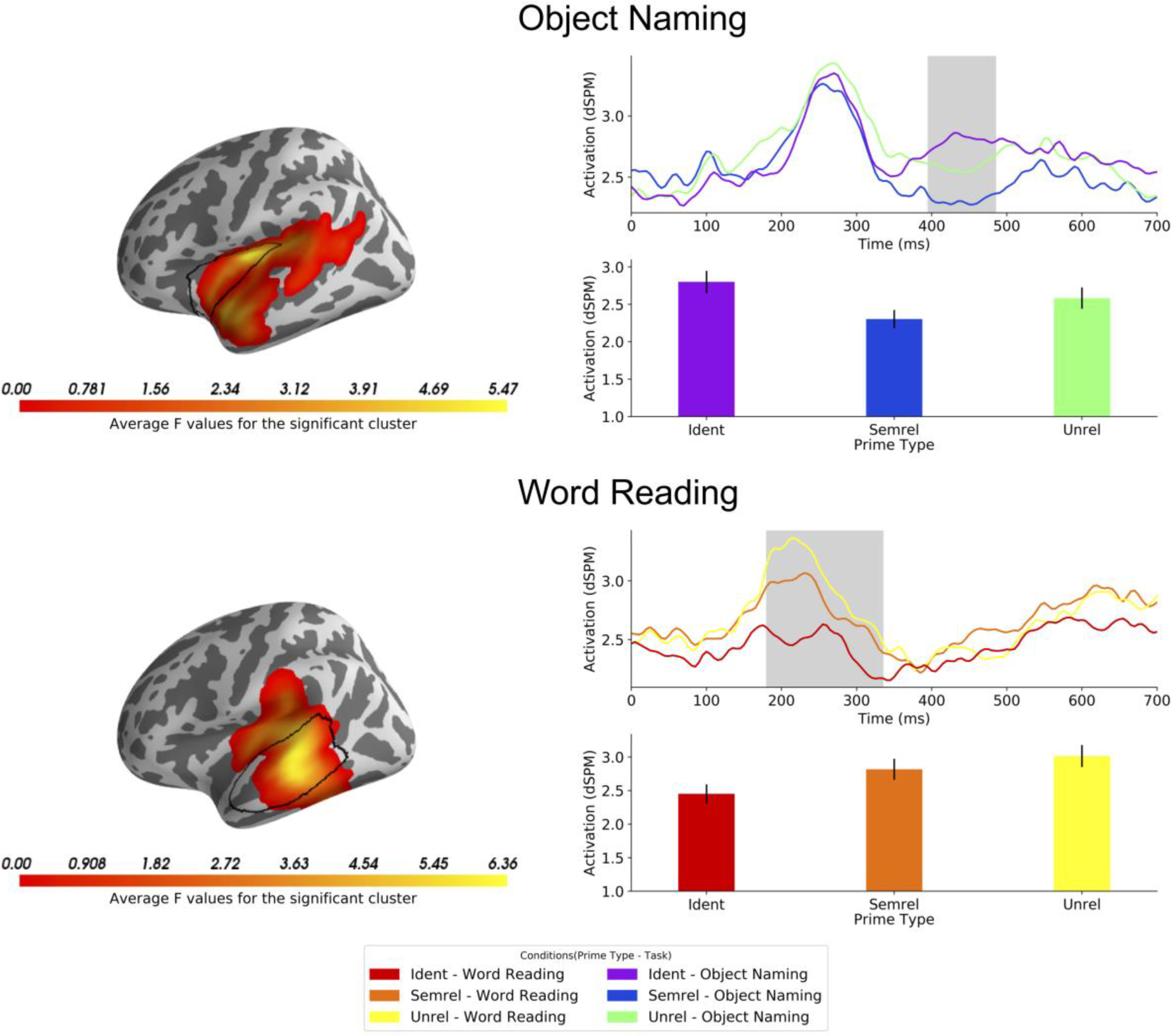
Within task analysis of priming effects, showing an amplitude reduction centered around insular cortex (borders indicated by the black line) for the semantically related condition in Object Naming (top) and a step-wise amplitude reduction (Ident < Semrel < Unrel) in the Word Reading task centered around the left STG (borders indicated by the black line) (*p* < 0.05, corr.).

## DISCUSSION

Here we took advantage of the spatiotemporal resolution of MEG to disambiguate the sources of two effects in lexical access research that have critically shaped our understanding of how words are accessed in comprehension vs. production. Specifically, the fact that semantic relatedness delays naming times in production has led to the hypothesis that in production, semantically related words compete with each other during conceptual preparation and/or lexical search. Here, however, we found no support for this hypothesis. Instead, our direct comparison of language comprehension and production revealed that while the semantic facilitory effect of comprehension localized in left middle temporal cortex in a time window consistent with lexical access, the inhibitory effect of production did not. Instead, a pattern consistent with inhibition was observed in left posterior insular cortex at 395-485ms after picture onset, well after lexical access in most models. This conforms to a motor preparation account of the production effect. In what follows we discuss the theoretical consequences and empirical limitations of these findings.

### Facilitation in word reading at 180-335ms centered in left superior temporal cortex

A classic semantic priming effect exhibits a reduction of amplitude as a function of semantic similarity between prime and target. Thus in the current paradigm, we expected facilitation to manifest as lower amplitudes for the semantically related condition than the unrelated condition and even lower ones for the identity condition, which involves repetition of the same word. Exactly this pattern was observed in left superior temporal cortex at 180-335 ms, which conforms with the localization of lexical access both in space (Hickok & Poeppel, 2007; Hillis, Rorden, & Fridriksson, 2017; Lau, Phillips, & Poeppel, 2008) and time (“lemma selection” in Indefrey, 2011; Indefrey & Levelt, 2004). Thus, this effect has a straightforward interpretation in terms of spreading activation between semantically related lexical representations. Research on reading aloud has also shown the STG to be connected to other areas relevant for semantic processing, such as the angular gyrus and the inferior temporal sulcus, forming a broader semantic network for reading aloud (Boukrina & Graves, 2013). Our positive priming effect in word reading formed a clear comparison point for the effect of semantic relatedness in object naming.

### Interference in object naming at 395-485ms centered in and around left insular cortex

In contrast to word reading, our MEG data for picture naming showed a pattern in which the semantically related condition separated both from the unrelated and the identity conditions. This pattern is consistent with similarity-based interference between related, but not identical, meanings. Crucially, the time resolution of our measurement allowed us to determine whether a possible interference pattern occurs early at 100-300ms, during conceptual preparation or lexical access, or later, at the stage of motor preparation. Our results clearly conformed to the latter hypothesis, showing a significant interference pattern at 395-485ms, centering in left insular cortex, a region that has been linked to motor planning of articulation (Ackermann & Riecker, 2004; Dronkers, 1996; Wise et al., 1999), and spreading to anterior parts of the STG. Crucially, in classic models of object naming, the timing of the effect coincides with the estimated timings of phonetic encoding and articulatory planning (Indefrey, 2011; Indefrey & Levelt, 2004).

As regards mechanisms, our results are consistent with the so-called Response Exclusion Hypothesis (Janssen et al., 2008; Mahon et al., 2007), in which the interference effect is generated at the motor output buffer. A core principle of this argument is that words have a privileged access to the motor preparation system over images. As a result, the primes have to be excluded from the single-channel motor preparation buffer before the target image can be named, which becomes more difficult as the prime and the target become more semantically related. This hypothesis may also provide a way to think about the directionality of our effect, which manifested as an amplitude reduction for semantically related targets as compared to the other two conditions: the larger amplitudes of the unrelated and identity conditions may reflect a process of more synchronous motor preparation for these conditions.

Importantly, the crucial aspect of our findings lies in the temporal information. The 395-485ms cluster observed for the interference effect occurred significantly later than conceptual preparation and lexical access stages, as depicted by models of object naming (Indefrey, 2011; Indefrey & Levelt, 2004). Thus, given the data observed here, we find no support for the hypothesis positing an early conceptual locus for interference, and instead, our findings favor the hypothesis of later motor level interference. The spatial clusters that we found complement these conclusions showing centralized activity at the STG associated with the facilitory effect in comprehension, and around the insular cortex for the interference effect in production.

### Widespread effect of task

We observed a widespread effect of task that covered almost the entire left hemisphere, throughout the whole analysis time window. This might initially seem surprising considering our attempt to design our tasks to be maximally similar. However, this large effect might be driven not only by the contrasting tasks, but also by the contrasting modalities of the primes. In fact, in Word Reading, the prime and the target are both written words, whereas in Object Naming, the prime is word while the target is an image. Therefore, our widespread effect could have been driven by the tasks themselves, or by matching vs. mismatching modalities of the prime and target. We speculate that the effect was likely driven by both of these contrasts, given its extensive coverage in both time and space.

## CONCLUSION

This work shows evidence that semantic interference in production and facilitation in comprehension localize differently, both in space and time, with an early facilitation in the STG in comprehension and a late interference around the insular cortex in production. These findings challenge the hypothesis of an early conceptual locus of interference. The effect of SOA failed to interact with our other manipulations, indicating that the range of SOAs that we used does not seem to have an impact on priming effects. Essentially, our findings suggest that while the early stages of comprehension involve co-activation of multiple related meanings, such co-activation may be absent in production.

## Acknowledgments

This work was supported by the New York University Abu Dhabi Institute (Grant G1001).

